# Brain Region-Specific Gene Co-expression Networks Reveal Neuroinflammation and ER Stress Signatures in Major Depressive Disorder

**DOI:** 10.1101/2025.06.18.660280

**Authors:** Lejun Li, Meiqi Wang, Yanbing Jie

## Abstract

This study aims to investigate gene expression changes in different brain regions of Major Depressive Disorder (MDD) patients using transcriptomics and bioinformatics approaches. Through differential expression gene (DEG) analysis, functional enrichment analysis, and Weighted Gene Co-expression Network Analysis (WGCNA), we aimed to identify key genes, pathways, and co-expression modules associated with MDD, with particular emphasis on the role of neuroinflammation and endoplasmic reticulum (ER) stress pathways in MDD pathology and their region-specific characteristics. The findings of this study will provide novel insights into the complex molecular mechanisms of MDD and lay the foundation for the future development of diagnostic biomarkers and therapeutic targets.

## Introduction

Major Depressive Disorder (MDD) is a debilitating psychiatric illness characterized by severe mood disturbances, anhedonia, and cognitive dysfunction, affecting millions worldwide. Despite its significant public health burden, the precise molecular mechanisms underlying MDD remain largely elusive, hindering the development of effective and personalized treatments.[1] While traditional theories have focused on monoamine neurotransmitter imbalances, emerging evidence increasingly implicates a broader array of biological processes, including neuroinflammation and endoplasmic reticulum (ER) stress, in the pathophysiology of various central nervous system (CNS) disorders, including neuropsychiatric conditions.

Neuroinflammation, characterized by the activation of glial cells (e.g., microglia and astrocytes) and the release of pro-inflammatory cytokines, can lead to neuronal damage, synaptic dysfunction, and altered neurocircuitry, all of which are pertinent to MDD. Simultaneously, the endoplasmic reticulum, a crucial organelle for protein folding, modification, and transport, can experience stress when its capacity is overwhelmed, triggering the Unfolded Protein Response (UPR). Chronic activation of ER stress and UPR has been consistently linked to neurodegenerative diseases and is gaining recognition as a contributor to psychiatric disorders. The complex interplay between neuroinflammation and ER stress, forming a vicious cycle of cellular dysfunction, is increasingly recognized as a critical factor in brain pathology. However, the specific gene expression signatures of these interconnected biological processes, particularly their regional specificity across different brain areas, have not been systematically investigated in MDD patients.

To address these gaps, this study leveraged advanced transcriptomics and bioinformatics approaches on publicly available gene expression datasets (GSE80655 and GSE102556) from the Gene Expression Omnibus (GEO) database. Our primary objective was to comprehensively explore gene expression alterations in distinct brain regions of MDD patients. By employing differential expression gene (DEG) analysis, Gene Ontology (GO) functional enrichment analysis, Gene Set Enrichment Analysis (GSEA), and Weighted Gene Co-expression Network Analysis (WGCNA), we aimed to identify key genes, pathways, and co-expression modules associated with MDD. A particular focus was placed on elucidating the role of neuroinflammation and ER stress pathways in MDD pathology, as well as their brain region-specific characteristics. The findings from this research are expected to provide novel insights into the complex molecular mechanisms of MDD, identify potential diagnostic biomarkers, and suggest new avenues for therapeutic intervention.

## Methods

This study aimed to deeply investigate gene expression changes in various brain regions of Major Depressive Disorder (MDD) patients and their associations with neuroinflammation and endoplasmic reticulum (ER) stress pathways. The research workflow encompassed differential expression gene analysis, functional enrichment analysis, and Weighted Gene Co-expression Network Analysis.

### Sample and Data Processing

This study utilized publicly available gene expression data from the Gene Expression Omnibus (GEO) database, specifically datasets GSE80655 and GSE102556. These datasets contain transcriptomic information from MDD patients and healthy control subjects across different brain regions. Raw count data underwent quality control prior to subsequent bioinformatics analyses.[2][3][4][5]

### Differential Expression Gene Analysis

Differential Expressed Genes (DEGs) analysis was performed using the DESeq2 R package. Raw count data were first subjected to normalization to mitigate batch effects. Subsequently, a generalized linear model was employed to identify genes with significantly differential expression between MDD patients and healthy controls. To control for false positives arising from multiple hypothesis testing, the Benjamini-Hochberg method was applied for multiple testing correction. Finally, genes with an absolute log2FoldChange (|log2FoldChange|) > 1 and a Benjamini-Hochberg adjusted P-value < 0.05 were considered as differentially expressed.[6]

### Functional Enrichment Analysis

Functional enrichment analysis was conducted using the clusterProfiler R package to perform Gene Ontology (GO) functional enrichment analysis on differentially expressed genes, with a primary focus on Biological Process terms. This analysis aimed to identify the enrichment of DEGs in specific biological functions, thereby elucidating biological pathways potentially affected in MDD patients. The significance of enrichment was assessed using a hypergeometric test, with an adjusted P-value < 0.05 as the criterion for significant enrichment. Enrichment results were visualized using dot plots and bar plots, offering an intuitive representation of the number of enriched pathways, their significance, and associated genes.[7]

### Weighted Gene Co-expression Network Analysis

Weighted Gene Co-expression Network Analysis (WGCNA)[8] was performed using the WGCNA R package. The aim was to construct gene co-expression networks and identify gene modules exhibiting similar expression patterns, further exploring the association of these modules with MDD phenotypes. Initially, gene expression data underwent quality control: genes with a standard deviation less than 0.5 were removed to exclude those with minimal expression variation and limited information content. Subsequently, sample clustering was examined to identify and remove potential outlier samples, ensuring data quality.

To construct a scale-free network, the optimal soft thresholding parameter (soft power, β) was selected by analyzing the scale-free topology of the network across different soft thresholds. In this study, a soft threshold β = 14 was chosen, as this parameter ensured the network met the scale-free topology criterion (scale-free topology R^2^ > 0.8), thereby ensuring the biological plausibility of the network. Gene co-expression modules were identified using the dynamic tree cut method, with a minimum module size of 60 genes to ensure each module possessed sufficient statistical power and biological interpretability. To merge highly correlated and similar modules, the correlation between module eigengenes was calculated, and modules with high correlation were merged, with a merging threshold set at 0.25.

Functional enrichment analysis was performed on the identified gene modules using GO databases[9][10] to annotate their biological functions, thereby elucidating the biological processes represented by each gene module. Module eigengenes were utilized to assess the overall expression pattern of each module.

## Results

This study conducted an in-depth analysis of gene expression profiles in different brain regions of MDD patients, revealing molecular characteristics associated with neuroinflammation, ER stress, and related cellular damage and neurological dysfunction in MDD.

### Differential Expression Genes and Regional Functional Enrichment Analysis

Differential expression gene analysis using DESeq2 identified multiple sets of significantly differentially expressed genes between MDD patients and healthy control subjects. We performed Gene Ontology (GO) Biological Process enrichment analysis for differentially expressed genes identified in the Cg25, nAcc, and Sub brain regions. The results showed that DEGs in these brain regions were significantly enriched in various stress responses (e.g., response to corticosterone, response to mineralocorticoid) and immune-related processes (e.g., humoral immune response, antimicrobial humoral immune response mediated by antimicrobial peptide), suggesting a pervasive stress and inflammatory response in the brains of MDD patients (Figure 1). Furthermore, the Cg25 region also showed enrichment in the regulation of nerve impulse transmission, the nAcc region in pathways related to musculoskeletal development, while the Sub region uniquely enriched for cilium assembly and neurotransmitter uptake (e.g., dopamine uptake, catecholamine uptake), indicating certain functional specific alterations across brain regions.

**FIGURE 1.**
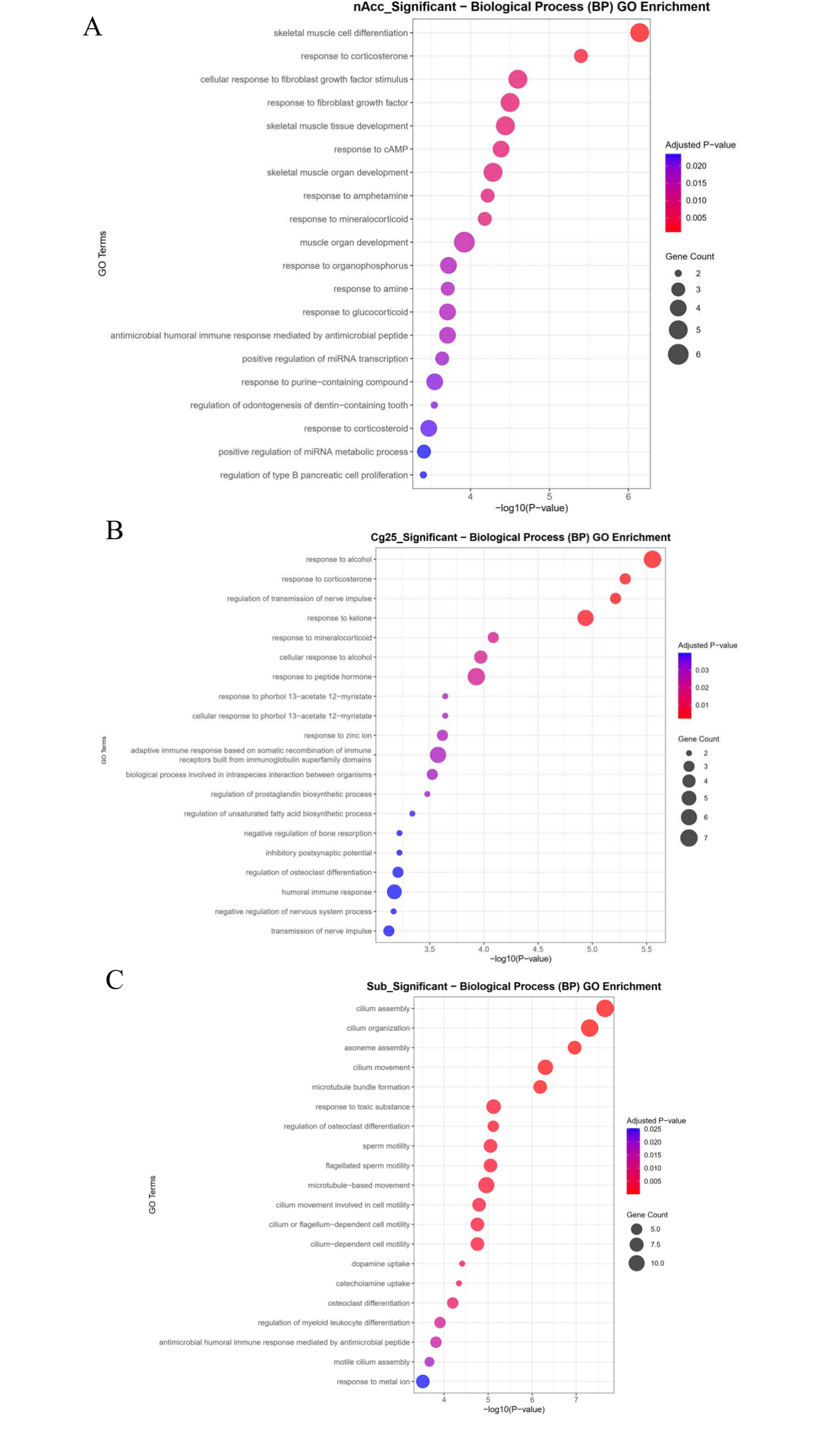

### WGCNA Identifies Core MDD-Related Gene Co-expression Modules

Weighted Gene Co-expression Network Analysis (WGCNA) identified multiple gene co-expression modules. Among them, the module eigengene score of the turquoise module showed a certain difference between MDD patient samples and the normal control group (MDD group average value was 0.21, control group average value was -0.21), suggesting that the overall expression pattern of genes in this module changed in MDD. GO enrichment analysis of genes within the turquoise module showed that this module was significantly enriched in terms related to cell adhesion, extracellular matrix organization, gliogenesis and glial cell differentiation, suggesting abnormal activity of glial cells and remodeling of the brain microenvironment. At the cellular component level, structures related to intracellular homeostasis and autophagy, such as vacuolar membrane and lysosomal membrane, were significantly enriched. At the molecular function level, ion channel activity (including voltage-gated and metal ion transmembrane transport), extracellular matrix structural constituent, and growth factor binding were also significantly enriched (Figure 2). These findings collectively depict that the turquoise module in MDD may participate in neuroinflammation, stress response, and neural circuit dysfunction by regulating glial cell function, intracellular membrane system homeostasis, and neuronal electrophysiological activity.

**FIGURE 2.**
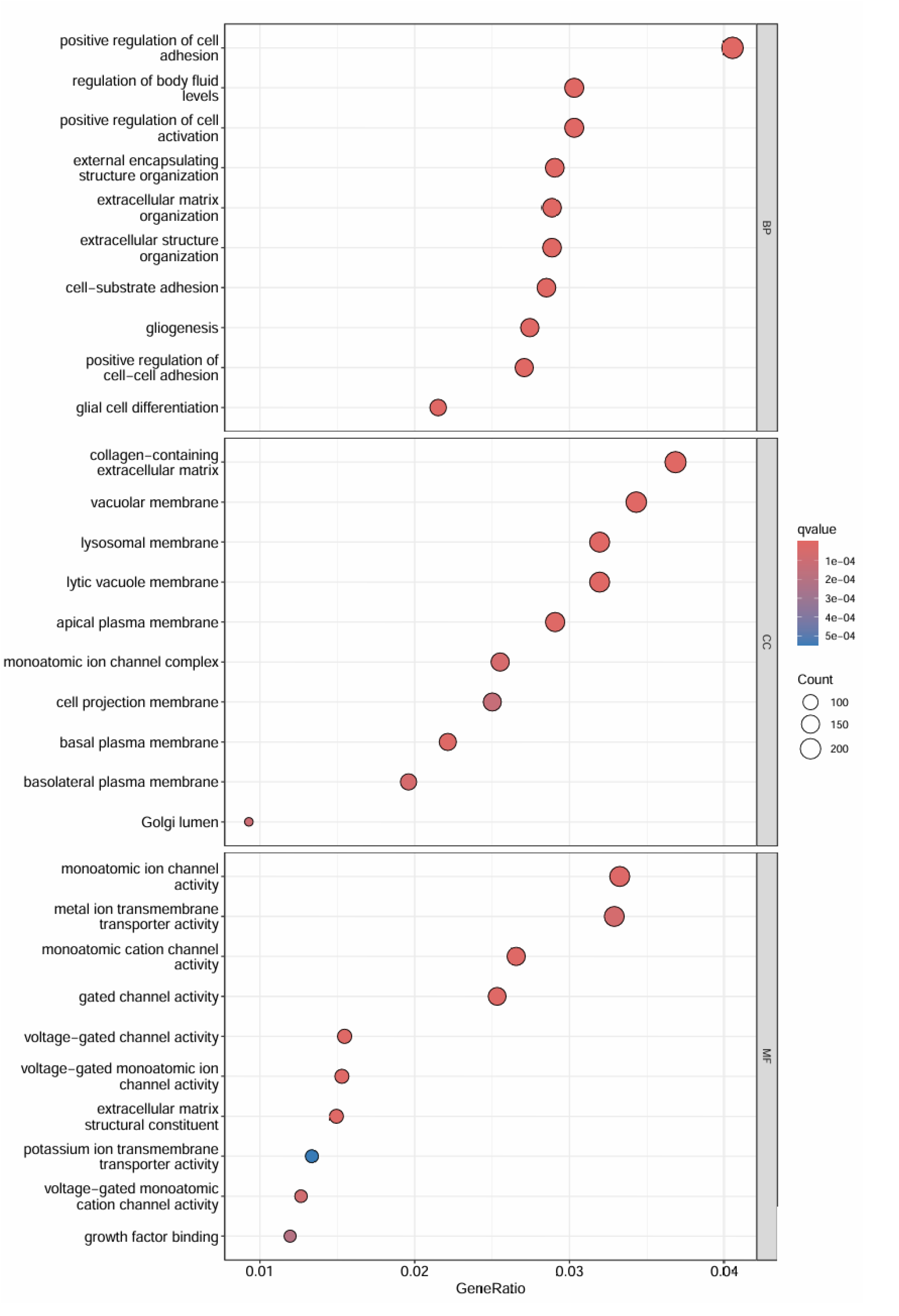

### Regional GSEA Reveals Widespread Activation of Neuroinflammation and ER Stress Pathways

To further delve into the regional specificity of MDD-related pathways, we performed GSEA enrichment analysis on the Cg25, nAcc, and Sub brain regions. GSEA enrichment score curves showed that multiple pathway gene sets exhibited positive enrichment in these brain regions in MDD patients. GSEA dot plots provided more detailed pathway enrichment information (Figure 3):

**FIGURE 3.**
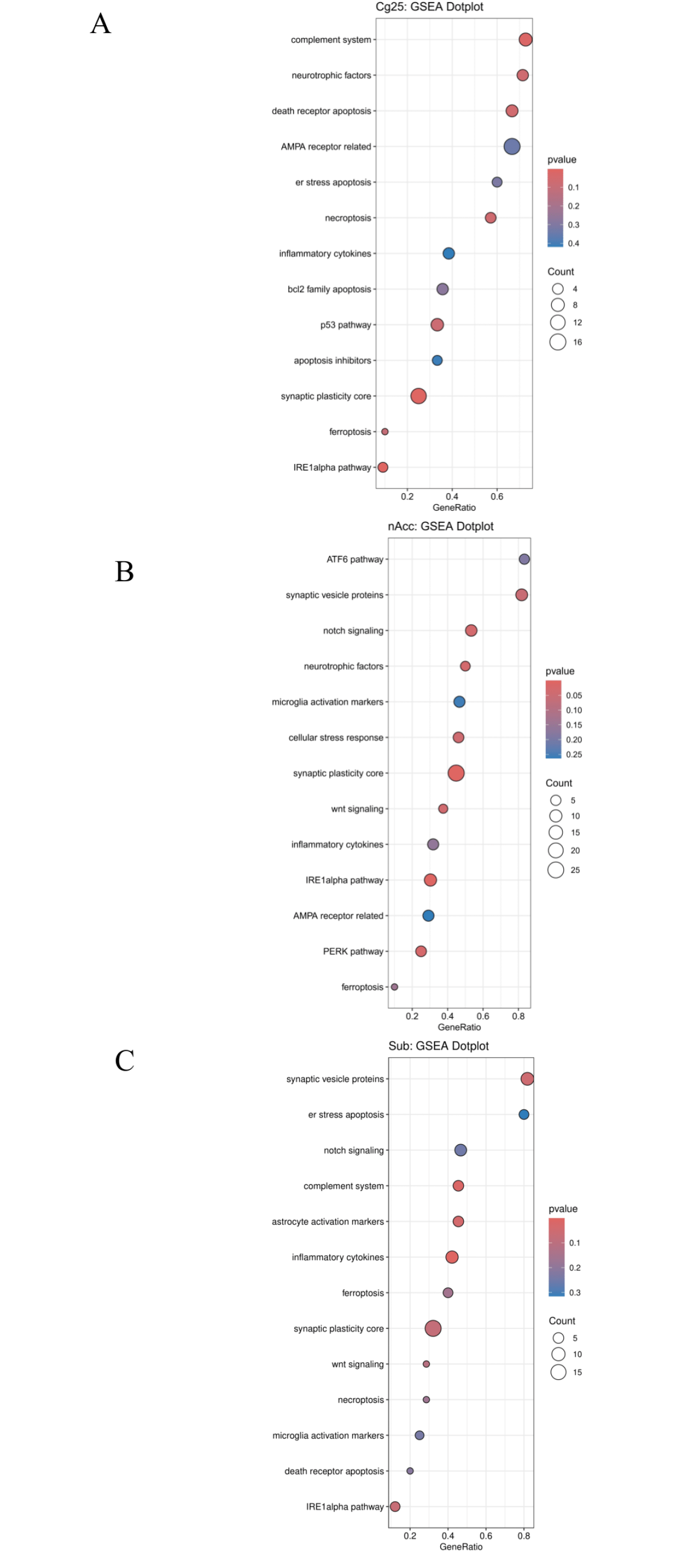

- **Cg25 brain region:** Significantly enriched for inflammatory pathways such as “inflammatory cytokines” and “complement system.” More importantly, the enrichment of “ER stress apoptosis,” “IRE1alpha pathway,” and various cell death pathways (e.g., ferroptosis, necroptosis) provided direct evidence of ER stress activation and widespread cell death in the Cg25 brain region. Additionally, the enrichment of “neurotrophic factors” and “synaptic plasticity core” also suggested potential impairment in neural function and plasticity.
- **nAcc brain region:** Showed the most significant and comprehensive ER stress activation features, with all three core branches of UPR:”ATF6 pathway,” “IRE1alpha pathway,” and “PERK pathway”, being significantly enriched. Simultaneously, the enrichment of “microglia activation markers” and “inflammatory cytokines” directly confirmed neuroinflammatory activation in the nAcc brain region. Furthermore, neuro-functional related pathways such as “synaptic vesicle proteins” and “synaptic plasticity core” were also significantly upregulated, indicating alterations in synaptic integrity and function.
- **Sub brain region:** GSEA results showed the enrichment of “astrocyte activation markers,” “microglia activation markers,” “inflammatory cytokines,” and “complement system,” clearly indicating neuroglia-mediated neuroinflammation in the Sub brain region. Concurrently, the enrichment of “ER stress apoptosis,” “IRE1alpha pathway,” and various cell death pathways (e.g., ferroptosis, necroptosis) again emphasized the active ER stress and cellular damage in this region.

### Integration of Key Modules and Regional Findings

The WGCNA and regional GSEA results of this study are highly consistent and mutually corroborative. The turquoise module identified by WGCNA not only showed overall differences in MDD, but its genes were also significantly enriched in inflammation, UPR, and depression-related pathways. This module had a higher module eigengene score in the nucleus accumbens (nAcc) brain region (0.13). The GSEA results for the nAcc brain region also provided strong molecular evidence, indicating the simultaneous activation of the three core branches of UPR, neuroglia-mediated inflammation, and pathways related to neural function in this region. These findings collectively depict a complex molecular landscape of neuroinflammation, ER stress, and related cellular damage and neurological dysfunction in MDD, and suggest that nAcc may be a core brain region connecting these pathological processes.

## Supporting information

manuscript

Deborah A Siegele, Rex L Chisholm, Petra Fey, Maria Cristina Aspromonte, Maria Victoria Nugnes, Federica Quaglia, Silvio Tosatto, Michelle Giglio, Suvarna Nadendla, Giulia Antonazzo, Helen Attrill, Gil dos Santos, Steven Marygold, Victor Strelets, Christopher J Tabone, Jim Thurmond, Pinglei Zhou, Saadullah H Ahmed, Praoparn Asanitthong, Diana Luna Buitrago, Meltem N Erdol, Matthew C Gage, Mohamed Ali Kadhum, Kan Yan Chloe Li, Miao Long, Aleksandra Michalak, Angeline Pesala, Armalya Pritazahra, Shirin C C Saverimuttu, Renzhi Su, Kate E Thurlow, Ruth C Lovering, Colin Logie, Snezhana Oliferenko, Judith Blake, Karen Christie, Lori Corbani, Mary E Dolan, Harold J Drabkin, David P Hill, Li Ni, Dmitry Sitnikov, Cynthia Smith, Alayne Cuzick, James Seager, Laurel Cooper, Justin Elser, Pankaj Jaiswal, Parul Gupta, Pankaj Jaiswal, Sushma Naithani, Manuel Lera-Ramirez, Kim Rutherford, Valerie Wood, Jeffrey L De Pons, Melinda R Dwinell, G Thomas Hayman, Mary L Kaldunski, Anne E Kwitek, Stanley J F Laulederkind, Marek A Tutaj, Mahima Vedi, Shur-Jen Wang, Peter D’Eustachio, Lucila Aimo, Kristian Axelsen, Alan Bridge, Nevila Hyka-Nouspikel, Anne Morgat, Suzi A Aleksander, J Michael Cherry, Stacia R Engel, Kalpana Karra, Stuart R Miyasato, Robert S Nash, Marek S Skrzypek, Shuai Weng, Edith D Wong, Erika Bakker, Tanya Z Berardini, Leonore Reiser, Andrea Auchincloss, Kristian Axelsen, Ghislaine Argoud-Puy, Marie-Claude Blatter, Emmanuel Boutet, Lionel Breuza, Alan Bridge, Cristina Casals-Casas, Elisabeth Coudert, Anne Estreicher, Maria Livia Famiglietti, Marc Feuermann, Arnaud Gos, Nadine Gruaz-Gumowski, Chantal Hulo, Nevila Hyka-Nouspikel, Florence Jungo, Philippe Le Mercier, Damien Lieberherr, Patrick Masson, Anne Morgat, Ivo Pedruzzi, Lucille Pourcel, Sylvain Poux, Catherine Rivoire, Shyamala Sundaram, Alex Bateman, Emily Bowler-Barnett, Hema Bye-A-Jee, Paul Denny, Alexandr Ignatchenko, Rizwan Ishtiaq, Antonia Lock, Yvonne Lussi, Michele Magrane, Maria J Martin, Sandra Orchard, Pedro Raposo, Elena Speretta, Nidhi Tyagi, Kate Warner, Rossana Zaru, Alexander D Diehl, Raymond Lee, Juancarlos Chan, Stavros Diamantakis, Daniela Raciti, Magdalena Zarowiecki, Malcolm Fisher, Christina James-Zorn, Virgilio Ponferrada, Aaron Zorn, Sridhar Ramachandran, Leyla Ruzicka, Monte Westerfield, The Gene Ontology knowledgebase in 2023, *Genetics*, Volume 224, Issue 1, May 2023, iyad031, https://doi.org/10.1093/genetics/iyad031

## Reference

1. Karrouri R, Hammani Z, Benjelloun R, Otheman Y. Major depressive disorder: Validated treatments and future challenges. World J Clin Cases. 2021 Nov 6;9(31):9350–9367. doi: 10.12998/wjcc.v9.i31.9350. PMID: 34877271; PMCID: PMC8610877.

2. Edgar R, Domrachev M, Lash AE. Gene Expression Omnibus: NCBI gene expression and hybridization array data repository Nucleic Acids Res. 2002 Jan 1;30(1):207–10

3. Ramaker RC, Bowling KM, Lasseigne BN, Hagenauer MH et al. Post-mortem molecular profiling of three psychiatric disorders. Genome Med 2017 Jul 28;9(1):72. PMID: 28754123

4. Labonté B, Engmann O, Purushothaman I, Menard C et al. Sex-specific transcriptional signatures in human depression. Nat Med 2017 Sep;23(9):1102-1111. PMID: 28825715

5. Mansouri S, Pessoni AM, Marroquín-Rivera A, Parise EM et al. Transcriptional dissection of symptomatic profiles across the brain of men and women with depression. Nat Commun 2023 Oct 26;14(1):6835. PMID: 37884562

6. Love MI, Huber W, Anders S. Moderated estimation of fold change and dispersion for RNA-seq data with DESeq2. Genome Biol. 2014;15(12):550. doi: 10.1186/s13059-014-0550-8. PMID: 25516281; PMCID: PMC4302049.

7. Yu G, Wang LG, Han Y, He QY. clusterProfiler: an R package for comparing biological themes among gene clusters. OMICS. 2012 May;16(5):284–7. doi: 10.1089/omi.2011.0118. Epub 2012 Mar 28. PMID: 22455463; PMCID: PMC3339379.

8. Langfelder, P., Horvath, S. WGCNA: an R package for weighted correlation network analysis. BMC Bioinformatics 9, 559 (2008). 10.1186/1471-2105-9-559

9. Ashburner M, Ball CA, Blake JA, Botstein D, Butler H, Cherry JM, Davis AP, Dolinski K, Dwight SS, Eppig JT, Harris MA, Hill DP, Issel-Tarver L, Kasarskis A, Lewis S, Matese JC, Richardson JE, Ringwald M, Rubin GM, Sherlock G. Gene ontology: tool for the unification of biology. The Gene Ontology Consortium. Nat Genet. 2000 May;25(1):25–9. doi: 10.1038/75556. PMID: 10802651; PMCID: PMC3037419.

10. The Gene Ontology Consortium, Suzi A Aleksander, James Balhoff, Seth Carbon, J Michael Cherry, Harold J Drabkin, Dustin Ebert, Marc Feuermann, Pascale Gaudet, Nomi L Harris, David P Hill, Raymond Lee, Huaiyu Mi, Sierra Moxon, Christopher J Mungall, Anushya Muruganugan, Tremayne Mushayahama, Paul W Sternberg, Paul D Thomas, Kimberly Van Auken, Jolene Ramsey, Deborah A Siegele, Rex L Chisholm, Petra Fey, Maria Cristina Aspromonte, Maria Victoria Nugnes, Federica Quaglia, Silvio Tosatto, Michelle Giglio, Suvarna Nadendla, Giulia Antonazzo, Helen Attrill, Gil dos Santos, Steven Marygold, Victor Strelets, Christopher J Tabone, Jim Thurmond, Pinglei Zhou, Saadullah H Ahmed, Praoparn Asanitthong, Diana Luna Buitrago, Meltem N Erdol, Matthew C Gage, Mohamed Ali Kadhum, Kan Yan Chloe Li, Miao Long, Aleksandra Michalak, Angeline Pesala, Armalya Pritazahra, Shirin C C Saverimuttu, Renzhi Su, Kate E Thurlow, Ruth C Lovering, Colin Logie, Snezhana Oliferenko, Judith Blake, Karen Christie, Lori Corbani, Mary E Dolan, Harold J Drabkin, David P Hill, Li Ni, Dmitry Sitnikov, Cynthia Smith, Alayne Cuzick, James Seager, Laurel Cooper, Justin Elser, Pankaj Jaiswal, Parul Gupta, Pankaj Jaiswal, Sushma Naithani, Manuel Lera-Ramirez, Kim Rutherford, Valerie Wood, Jeffrey L De Pons, Melinda R Dwinell, G Thomas Hayman, Mary L Kaldunski, Anne E Kwitek, Stanley J F Laulederkind, Marek A Tutaj, Mahima Vedi, Shur-Jen Wang, Peter D’Eustachio, Lucila Aimo, Kristian Axelsen, Alan Bridge, Nevila Hyka-Nouspikel, Anne Morgat, Suzi A Aleksander, J Michael Cherry, Stacia R Engel, Kalpana Karra, Stuart R Miyasato, Robert S Nash, Marek S Skrzypek, Shuai Weng, Edith D Wong, Erika Bakker, Tanya Z Berardini, Leonore Reiser, Andrea Auchincloss, Kristian Axelsen, Ghislaine Argoud-Puy, Marie-Claude Blatter, Emmanuel Boutet, Lionel Breuza, Alan Bridge, Cristina Casals-Casas, Elisabeth Coudert, Anne Estreicher, Maria Livia Famiglietti, Marc Feuermann, Arnaud Gos, Nadine Gruaz-Gumowski, Chantal Hulo, Nevila Hyka-Nouspikel, Florence Jungo, Philippe Le Mercier, Damien Lieberherr, Patrick Masson, Anne Morgat, Ivo Pedruzzi, Lucille Pourcel, Sylvain Poux, Catherine Rivoire, Shyamala Sundaram, Alex Bateman, Emily Bowler-Barnett, Hema Bye-A-Jee, Paul Denny, Alexandr Ignatchenko, Rizwan Ishtiaq, Antonia Lock, Yvonne Lussi, Michele Magrane, Maria J Martin, Sandra Orchard, Pedro Raposo, Elena Speretta, Nidhi Tyagi, Kate Warner, Rossana Zaru, Alexander D Diehl, Raymond Lee, Juancarlos Chan, Stavros Diamantakis, Daniela Raciti, Magdalena Zarowiecki, Malcolm Fisher, Christina James-Zorn, Virgilio Ponferrada, Aaron Zorn, Sridhar Ramachandran, Leyla Ruzicka, Monte Westerfield, The Gene Ontology knowledgebase in 2023, Genetics, Volume 224, Issue 1, May 2023, iyad031, 10.1093/genetics/iyad031

