## Supplementary material for "Brain Region-Specific Gene Co-expression Networks Reveal Neuroinflammation and ER Stress Signatures in Major Depressive Disorder": manuscript

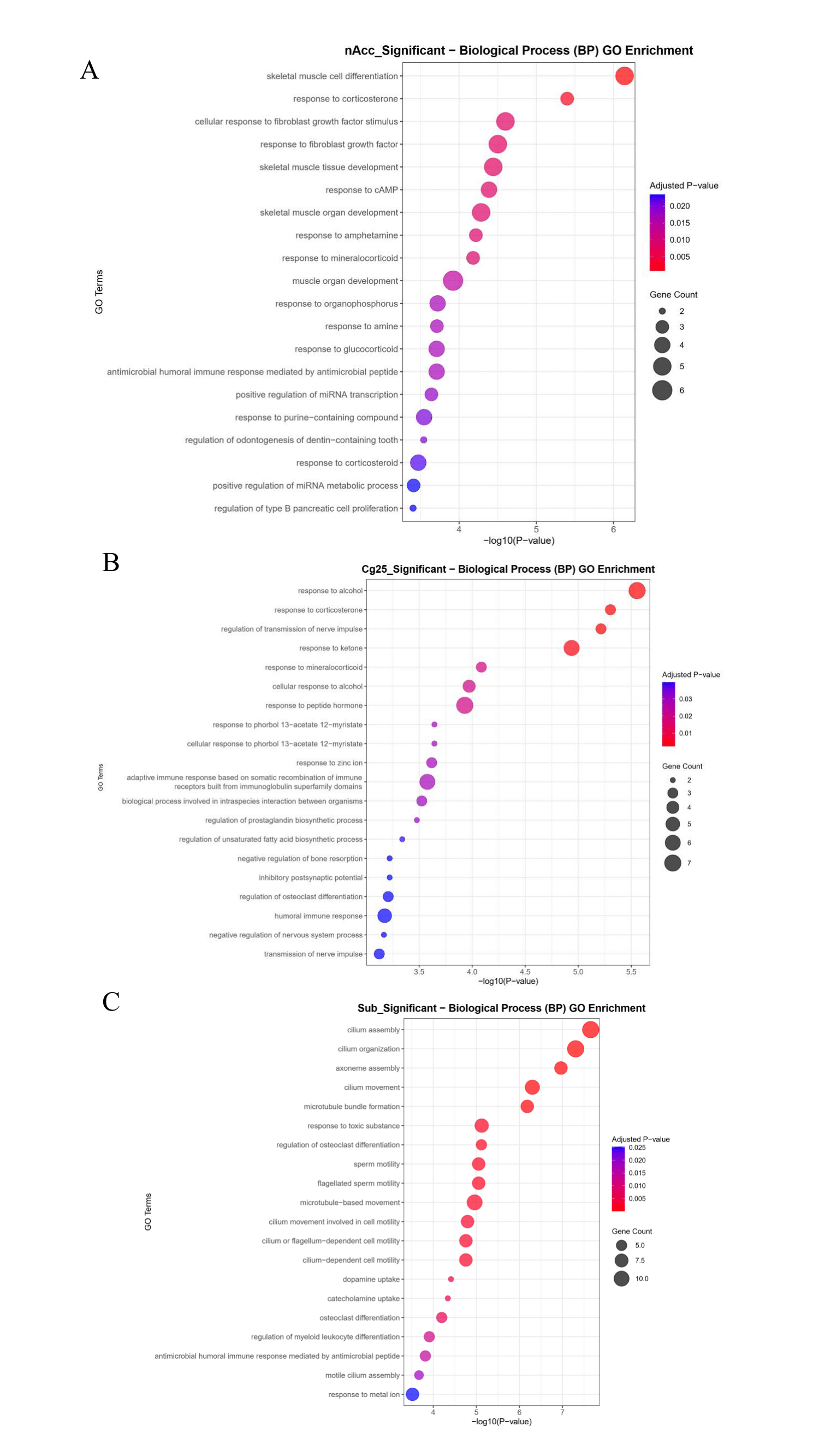

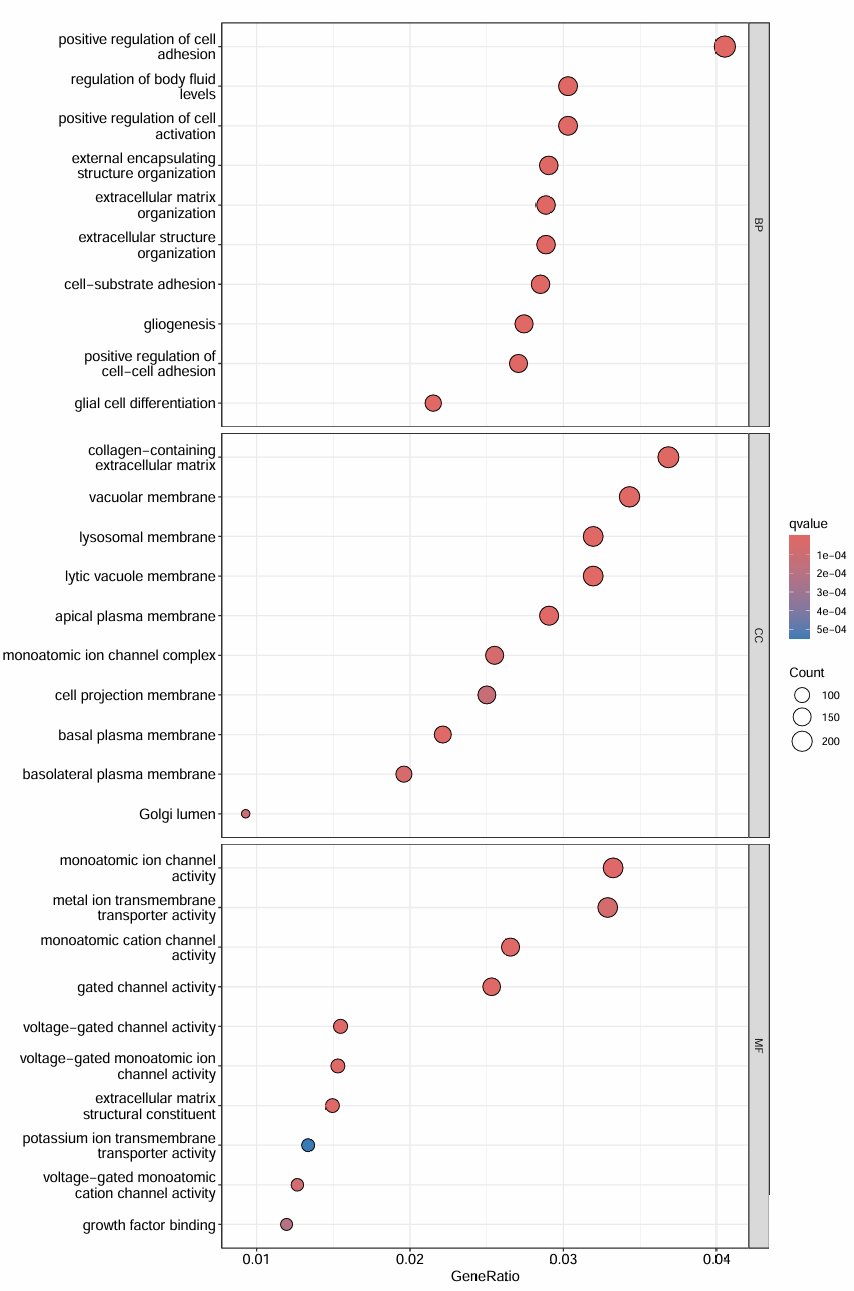

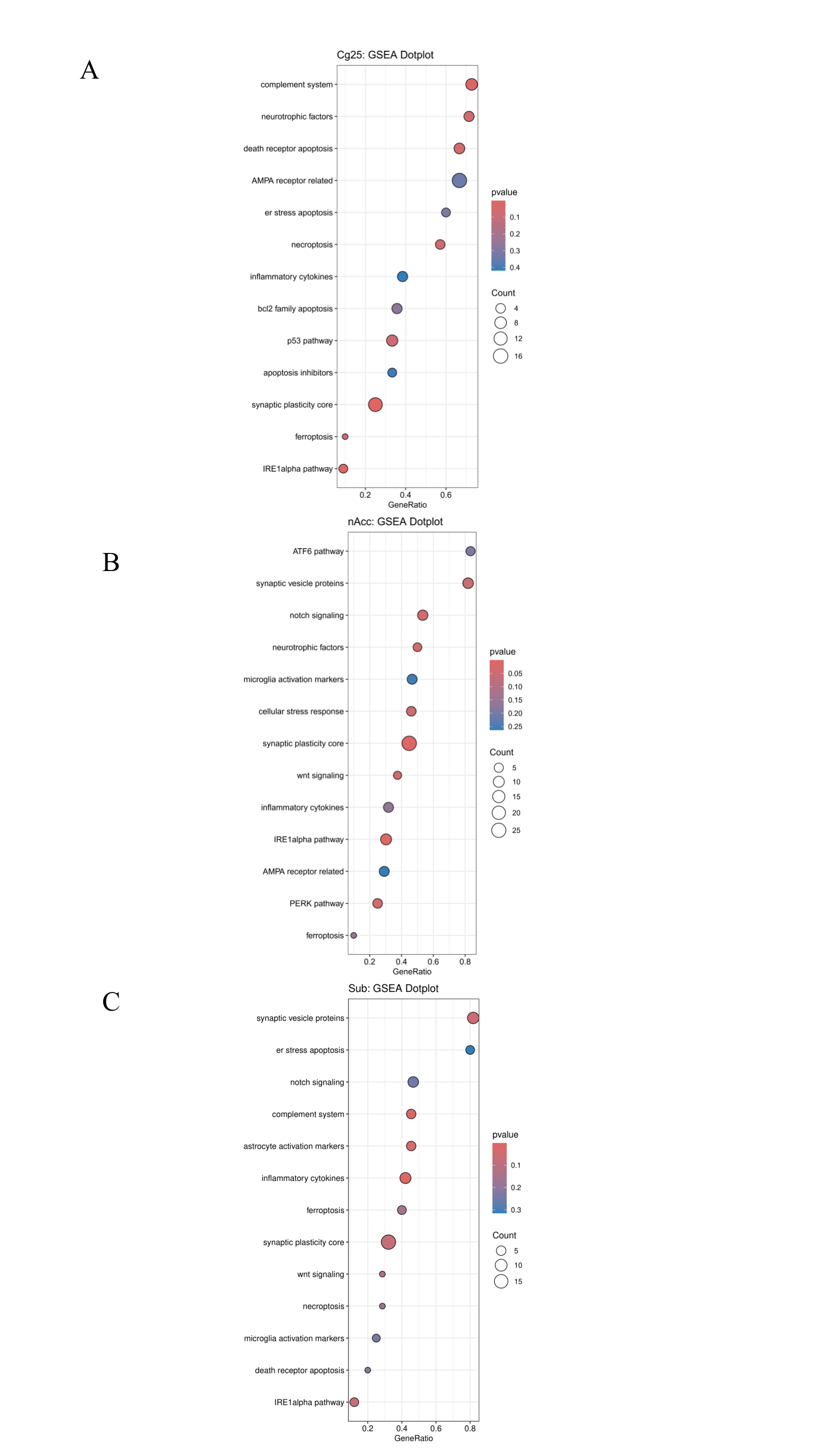
